# Measures of repetition suppression in the Fusiform Face Area are inflated by co-occurring effects of statistically learned visual associations

**DOI:** 10.1101/803163

**Authors:** Sophie-Marie Rostalski, Catarina Amado, Gyula Kovács, Daniel Feuerriegel

## Abstract

Repeated presentation of a stimulus leads to reductions in measures of neural responses. This phenomenon, termed repetition suppression (RS), has recently been conceptualized using models based on predictive coding, which describe RS as due to expectations that are weighted toward recently-seen stimuli. To evaluate these models, researchers have manipulated the likelihood of stimulus repetition within experiments. They have reported findings that are inconsistent across hemodynamic and electrophysiological measures, and difficult to interpret as clear support or refutation of predictive coding models. We instead investigated a different type of expectation effect that is apparent in stimulus repetition experiments: the difference in one’s ability to predict the identity of repeated, compared to unrepeated, stimuli. In previous experiments that presented pairs of repeated or alternating images, once participants had seen the first stimulus image in a pair, they could form specific expectations about the repeated stimulus image. However they could not form such expectations for the alternating image, which was often randomly chosen from a large stimulus set. To assess the contribution of stimulus predictability effects to previously observed RS, we measured BOLD signals while presenting pairs of repeated and alternating faces. This was done in contexts whereby stimuli in alternating trials were either i.) predictable through statistically learned associations between pairs of stimuli or ii.) chosen randomly and therefore unpredictable. We found that RS in the right FFA was much larger in trials with unpredictable compared to predictable alternating faces. This was primarily due to unpredictable alternating stimuli evoking larger BOLD signals than predictable alternating stimuli. We show that imbalances in stimulus predictability across repeated and alternating trials can greatly inflate measures of RS, or even mimic RS effects. Our findings also indicate that stimulus-specific expectations, as described by predictive coding models, may account for a sizeable portion of observed RS effects.

## 1. Introduction

Repeated presentation of a stimulus leads to reduced measures of neural responses, as observed using a variety of electrophysiological and neuroimaging techniques (for a review see Grill-Spector et al., 2006). Such effects are commonly known as repetition suppression (RS) or adaptation. Similarly, the correct and fulfilled expectation of a forthcoming stimulus also leads to reduced responses when compared to unexpected or surprising stimuli, for several stimulus categories and measures (known as expectation suppression, ES; for a review see Summerfield & Egner, 2009).

Explanations of repetition-as well as expectation-related phenomena under the framework of predictive coding (Rao & Ballard, 1999) have gained traction in recent years. This is because, in contrast to several other neurobiologically-plausible models of RS (for review see Grill-Spector et al., 2006), predictive coding models describe mechanisms that can potentially account for observed RS effects, and also how RS might be modulated by processes related to perceptual expectations and attention (e.g., Eger, 2004; Murray & Wojciulik, 2004). Predictive coding models conceptualize RS as a reduction of prediction error signals, due to perceptual expectations that are weighted toward recently-encountered stimuli (e.g., Friston, 2005; Grotheer & Kovács, 2015; Auksztulewicz & Friston, 2016). Factors such as attention are hypothesized to modulate the precision of sensory predictions (Feldman & Friston, 2010), which in turn influence the extent of observed RS. Predictive coding models describe different mechanisms than those in recently-formulated local circuit models of RS (Dhruv et al., 2011; Kaliukhovich & Vogels, 2016; Solomon & Kohn, 2014). However, the notion of precision in predictive coding models allows us to test hypotheses about how attention and perceptual expectations affect RS, while specific hypotheses have (to our knowledge) not yet been derived for the abovementioned local circuit models.

Summerfield et al. (2008) was the first to provide empirical support for the predictive coding model by showing that neuroimaging measures of RS can be modulated by contextual factors, such as the probability of stimulus repetition. They presented pairs of faces in each trial and reported that BOLD signal differences between repeated and unrepeated stimuli (i.e., repetition effects) were larger in blocks with high (75%), compared to blocks with low (25%) repetition probability. This interaction involving repetition probability was replicated several times using faces (for a review see Grotheer et al., 2014), and also for other stimulus categories such as letters (Grotheer & Kovács, 2014) and other non-face objects (Kronbichler et al., 2018; Mayrhauser et al., 2014). Notably, this interaction has mostly been reported in studies using fMRI; when using electrophysiological measures researchers have found separable, non-interacting repetition and expectation effects (Feuerriegel et al., 2018a; Kaliukhovich & Vogels, 2014; Todorovic & de Lange, 2012; Vinken et al., 2018), with the exception of Summerfield et al. (2011, but see Feuerriegel et al., 2018a for an alternative explanation of this result).

When interpreting these findings, it is important to differentiate the neural mechanisms of RS from how RS is typically measured within an experiment (as a difference between a comparable repeated and unrepeated stimulus condition). In such experiments, any effect that will influence repeated and unrepeated stimulus-evoked responses in different ways will also contribute to the measured magnitude of RS, even if that effect is unrelated to the underlying processes responsible for RS (reviewed in Feuerriegel, 2016). In Summerfield et al. (2008) and similar experiments, participants could learn to expect stimulus repetitions in the 75% repetition blocks, whereby in the same block unrepeated stimulus trials were relatively rare and surprising. Conversely, in the 25% repetition blocks the unrepeated stimuli were instead expected, and the repeated stimuli relatively surprising. Accordingly, the observed RS by expectation interaction could actually be produced by additive effects of genuine RS and a distinct expectation related suppression effect (ES; Kaliukhovich & Vogels, 2011; Larsson & Smith, 2012), with expectations suppressing responses to either repeated or unrepeated stimuli in different block types.

More recent studies have used “cue” stimuli, whereby the first stimulus in each trial signals the probability of stimulus repetition, in order to distinguish between additive and interactive effects of ES and RS. Todorovic and de Lange (2012) presented pairs of auditory tones, which could either repeat or change within a trial. The pitch of the first tone predicted stimulus alternation or repetition with 75% probability. They reported that RS and ES, as indexed by magnetoencephalography (MEG), were separable and occurred at distinct time windows. In a similar design using face stimuli Grotheer and Kovács (2015) reported that effects RS and ES on BOLD signals did not interact, and were partly dissociable in the time course of their effects on the hemodynamic response. In a follow-up study Amado and colleagues (2016) added a ‘neutral’ condition, in which expectations were not weighted toward either repeated or alternating stimuli, to separately quantify effects of fulfilled expectations and surprise. They found that surprise had a much larger effect on BOLD signals than fulfilled expectations, and that this effect of surprise was apparent for alternating (but not repeated) stimulus conditions (see also e.g., Figure 2 in De Gardelle et al., 2013; Figure 2 in Larsson & Smith, 2012). This suggests that, instead of ES modulating repetition effects, RS might in fact inhibit surprise-related response enhancements, as found in a recent EEG study (Feuerriegel et al., 2018b). These results, along with the inconsistency of findings across fMRI and electrophysiological recording methods, do not provide clear evidence that expectations modulate RS in the way previously specified by predictive coding models.

Besides expectations relating to stimulus repetition probability, there is another type of expectation that is prevalent in studies of RS, and relevant for evaluating predictive coding models of repetition effects. There is evidence from single-cell recordings of non-human primates (Meyer & Olson, 2011) as well as human electrophysiological and neuroimaging experiments (Turk-Browne et al., 2009; Hall et al., 2018; Pajani et al., 2017; Feuerriegel et al., 2018a) indicating that associations are formed between images that are shown temporally close together, and this association modulates neural responses. The proposed underlying mechanism is that the observers learn about the transitional statistics or rules of the stimulation sequences, as humans do from early childhood onwards to learn about their environment (Fiser & Aslin, 2002; Romberg & Saffran, 2011). As a seminal example, Meyer and Olson (2011) trained macaques to associate originally unrelated images by presenting the same stimulus pairs over a prolonged time period. The animals learned that one leading image was always followed by a specific trailing image. In a subsequent session, single-neuron activity was recorded from inferotemporal cortex (IT) while the animals viewed stimulus pairs which were either previously associated or randomly paired. IT neurons exhibited higher firing rates following stimuli which violated previously learned transitional rules, compared to those that were associated with the previous image.

This type of statistically learned expectation is relevant to a large number of stimulus repetition designs that have been used in the past. In these designs, participants are presented with two stimuli in each trial, which may be of the same or different identities. In repetition trials the identity of the second stimulus can be predicted after seeing the first stimulus in the trial, however the alternating (unrepeated) stimulus is often randomly-chosen from a set of multiple stimuli, and is very difficult to predict with any certainty (Feuerriegel, 2016). This imbalance in predictability across repetition and alternation trials could theoretically inflate the magnitude of, or even produce, many previously observed RS effects. Pajani et al. (2017) investigated this using a design that manipulated the predictability of the alternating stimuli. They presented stimuli in repetition blocks, composed of 75% repetition and 25% alternation trials, and alternation blocks, with only a 25% portion of repetition trials. Crucially, in a third block type 25% of trials were repetitions and 75% were predictable alternations, whereby the second stimulus was repeatedly paired with the first stimulus during a prior training session. They observed large differences in the magnitude of repetition effects, apparently due to reductions in BOLD signals for predictable compared to unpredictable alternating faces. Further evidence for predictability effects came from a recent EEG study (Feuerriegel et al., 2018a), who used a similar blocked design with predictable and unpredictable alternating faces. In the so-called “AB” blocks in that experiment the second stimulus in each trial could either be the same image as the first (repetition trials), or a specific same-sex face (predictable alternation trials). In the “AX” blocks, however, the second stimulus could either be a repetition of the first one, or a same-gender face, selected randomly from a set of 23 stimuli (unpredictable alternation trials). Differences in event-related potential (ERP) repetition effect magnitudes across AB and AX blocks were found during multiple time windows post stimulus onset. Importantly, these differences in observed repetition effects were due to differences in ERP responses to alternating stimuli across block types, and no differences across AB and AX blocks were found for repeating stimuli.

Critically, this study did not equate the relative novelty of AB and AX alternating stimuli, as each individual face identity was presented many more times in the AB compared to AX conditions. Similarly, in Pajani et al. (2017) the predictable alternating stimuli were presented many more times during the experiment than the unpredictable alternating stimuli, which were trial-unique. Because of this, it is unclear whether the observed effects were primarily due to effects of stimulus predictability or stimulus novelty, both of would have similar hypothesised effects on neural responses (e.g. Feuerriegel, 2016; Mur et al., 2010; Xiang and Brown, 1998).

We used a similar design to investigate the interplay of stimulus repetition and prediction effects using fMRI, while controlling for the relative novelty of predictable and unpredictable alternating stimuli. The previously introduced conditions in Feuerriegel et al., (2018a) were adopted, including predictable (AB) and unpredictable (AX) alternating trials. RS was measured by comparing BOLD signals in trials with repeated and alternating stimulus pairs. Importantly, prior to the fMRI scanning session participants underwent 4 training sessions on consecutive days, during which they were presented with 6 predictable alternating face pairs (i.e. the first face of a pair was always followed by a specific same-sex face) to create specific face associations for the alternating trials. Because previous fMRI studies that presented face stimuli (Amado et al., 2016; Egner et al., 2010; Summerfield et al., 2008) found the most pronounced effects of stimulus repetition and perceptual expectations in the fusiform face area (FFA; Kanwisher et al., 1997) we focused our analyses on this region.

Our design allowed us to control for effects of stimulus novelty, enabling a more accurate estimate of stimulus predictability effects in repetition designs. This also allowed us to assess whether this type of expectation may account for a portion of previously observed RS effects. To foreshadow our results, we found that predictability does modulate BOLD responses in the FFA and acts primarily upon responses to alternating stimuli, replicating the patterns effects in Pajani et al. (2017) and Feuerriegel et al., (2018a). While our results support the notion of separable RS and predictability effects, they also indicate that, when predictability is confounded with stimulus repetition, as in a large number of existing studies, RS effects are likely to be inflated (or perhaps even caused) by this predictability confound.

## 2. Methods

### 2.1 Participants

Twenty-two volunteers participated in the study. All were informed about the procedure of the study and gave written consent for participation beforehand. One participant was excluded due to not completing the experiment while another participant’s data was partially lost due to technical issues related to the MRI scanner. The remaining 20 participants (3 males; 4 left-handed) were between 19 and 28 years of age (*M* = 21.9, *SD* = 2.53). All had normal or corrected-to-normal vision. The experiment was conducted in accordance with the guidelines of the Declaration of Helsinki, and with the approval of the ethics committee of the University of Jena.

### 2.2 Stimuli

We presented 12 images of upright female faces as stimuli. Pictures were cropped to show faces without hair or clothes, resized to 440 x 400 pixels, converted to greyscale and equated in average luminance (Fig. 1A). Stimuli were presented against a black background using Psychtoolbox v.3.0.14 (Brainard, 1997; Kleiner et al., 2007) in MATLAB 2014a (The Mathworks). For each participant six stimuli were allocated randomly to be presented in the AB and the remaining six in the AX conditions.

**Figure 1.**
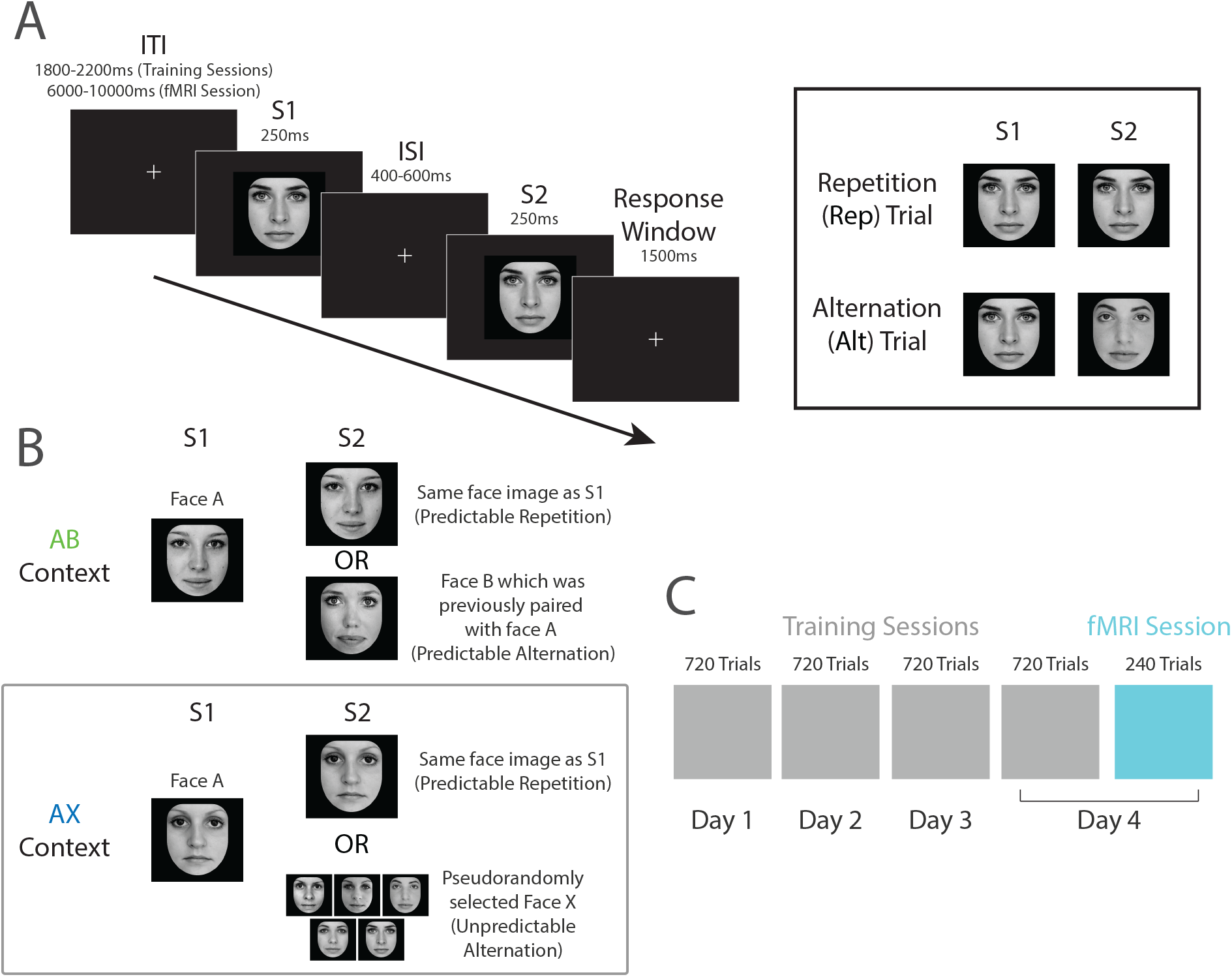
Trial structure and predictability cueing manipulation. A) In each trial S1 and S2 face stimuli were presented, separated by a 400-600 ms inter-stimulus interval (ISI). The S2 stimulus could either be the same face image as S1 (repetition trials) or a different female face (alternation trials). B) For alternation trials, the S2 face could either be a particular face “B” that was repeatedly paired with a specific S1 face “A” during the training sessions (AB context) or pseudorandomly-chosen from a set of 5 different faces (AX context). The probability of stimulus repetition was fixed at 50% across both contexts. C) Participants completed 4 training sessions over consecutive days. Trial structure, task (same-different forced choice), stimuli and AB/AX contexts were the same as in the fMRI scanning session but with a shorter ITI duration. Following the fourth training session participants then completed the fMRI session on the same day.

### 2.3 Experiment Design

Participants first completed a series of behavioural training sessions, followed by an fMRI session (see Fig. 1C). The experimental design, including the stimuli and task, was identical across training and fMRI data acquisition sessions, except where specified otherwise.

In each trial (Fig. 1A) an adapter (S1) and test stimulus (S2) were each presented for 250 ms, separated by an inter-stimulus interval (ISI) of 400-600 ms (randomised across trials). The image size of S2 was 20% smaller than that of S1 to avoid low-level adaptation processes. Trials were separated by an inter-trial interval (ITI): for the training sessions the ITI was 1800, 2000 or 2200 ms, randomly distributed across trials, and for the fMRI sessions it was 6, 8 or 10 seconds.

In each trial, S1 and S2 could either be identical (repetition trials; Rep) or depicting different identities (alternation trials; Alt). These trial types were presented in two different contexts (Fig. 1B), labelled as “AB” and “AX”. In the AB context the S2 face could either be a repetition of the S1 face (Rep trials), or a specific face identity that had previously been repeatedly paired and associated with the S1 identity during the training sessions (Alt trials). In these Alt trials of the AB context, each S1 face identity was consistently paired with one of the five other face identities that were allocated to the AB context. Each S1 identity in the AB stimulus set was paired with a different S2 face identity, ensuring that each face image would be presented an equal number of times throughout the experiment. In other words, once the participant has seen a given S1 face “A”, they could form expectations regarding the S2 to be a repetition of face “A” or a different, specific identity “B”. In the AX context S2 could either be the repetition of the S1 image, or a different identity, pseudo-randomly selected from the set of 5 other face identities. Therefore, in the AX context, there were no consistent pairings between S1 and S2 face identities for the Alt trials: S2 could be any of the five other faces, allocated to the AX condition, ensuring that each face appeared the same number of times throughout this condition. This procedure ensured further that each AB and AX face identity was presented the same number of times across the experiment. Thus altogether, we had two independent factors: trial type (Rep or Alt) and context, reflecting prior associations formed for Alt trials (AB) or not having such transitional rules (AX). The proportion of Rep trials (i.e., the probability of stimulus repetition) was 50% in both AB and AX contexts.

### 2.4 Procedure

Participants completed four training sessions across four consecutive days prior to the fMRI measurements. The fMRI session followed the last training session immediately on the fourth day. Each training session was composed of twelve blocks (60 trials per block, 720 trials per session) and lasted approximately 40 minutes. All sessions took place at approximately the same time of the day, in the afternoon hours to control for potential changes of attention that occur across the circadian cycle (Valdez et al., 2010).

During the training sessions participants learned the S1-S2 transition probabilities associated with face identities in the AB and AX contexts. Trials of AB and AX context were presented randomly interleaved within the same blocks of trials and with equal probability. For the AB context, each face image pairing (in Alt trials) was presented 120 times throughout the training sessions while for the AX context each of the possible Alt S1-S2 combinations was shown 24 times.

During the training and fMRI sessions the participants’ task was to decide whether the S1 and S2 were the same or different face images by pressing one of two keys on a keyboard (training session) or MRI-compatible button box (fMRI session). The spatial layout of response keys/buttons and associated response fingers were kept constant across behavioural and fMRI sessions. Instructions were presented in the centre of the screen prior to each run. Participants took a self-paced break between each run. The entire fMRI session lasted approximately 60 min.

### 2.5 Image Acquisition

Four experimental runs were completed, with each lasting for about 10 minutes and including 60 trials. A total of 240 trials were presented during the fMRI session. An additional localizer sequence was included to define the location of the FFA bilaterally (blocks of 40 images, size: 600 x 600 pixels on a grey background; exposition time: 300 ms, ISI: 200 ms; presenting faces, objects and Fourier-randomized noise patterns lasting for 20 seconds each). Using data from the localizer sequence we could identify the right FFA in 18 out of 20 participants (average MNI coordinates (± SE): 41 (1), −47 (1), −21 (1)). We could also define the left FFA in a subset of 14 participants (average MNI coordinates: −40 (1), −51 (2), −21 (1); p < 0.05 FWE) and included this ROI in a separate analysis.

Magnetic Resonance Images were acquired using a 3-Tesla magnetic resonance (MR) scanner from Siemens. For functional images, a standard T_2_-weighted echo-planar imaging (EPI) sequence (35 slices, 10° tilted relative to axial, TR = 2000 ms, echo time (TE) = 30 ms, flip angle 90°, 64 x 64 matrices, in plane resolution 3 mm isotopic voxel size) was used. A high resolution T_1_-weighted structural 3D scan was generated using a magnetization-prepared rapid gradient-echo (MP-RAGE; TR = 2300 ms; TE = 3.03 ms; 1 mm isotropic voxel size). For details of pre-processing and statistical analysis see Cziraki et al., (2010). Briefly, the functional images were realigned, normalized to the MNI-152 space, resampled to 2×2×2 mm resolution and spatially smoothed with a Gaussian kernel of 8 mm FWHM (SPM12, Welcome Department of Imaging Neuroscience, London, UK). A generalised linear model was specified, using the different conditions, as well as six movement parameters. Data from the localizer sequence were used to identify the location of the FFA individually by contrasting faces with objects and noise. From these coordinates the BOLD signal due to the experimental conditions was extracted using a 2 mm radius sphere, and the peak values were entered into the statistical models.

### 2.6 Statistical Analyses

Data and code required to reproduce all analyses will be available at https://osf.io/akygb/ at the time of publication. Statistical analyses were performed using Statistica (StatSoft) and JASP v0.9.1 (JASP Team). Mean response times and accuracy rates during the training sessions were analysed using 4 x 2 x 2 repeated measures ANOVAs with the factors of session (1, 2, 3, 4), context (AB, AX) and trial type (Rep, Alt). Peak BOLD signal values were analysed using a 2 x 2 repeated measures ANOVA with the factors context (AB, AX) and trial type (Rep, Alt). For all ANOVA models, Greenhouse-Geisser corrections were applied in cases where Mauchly tests indicated violations of sphericity. Additionally, we analysed the mean response times and accuracy rates for the fMRI session.

## 3. Results

### 3.1 Behavioral Results

A significant main effect of session was found for response times. Because the Mauchly test of sphericity revealed unequal variances of differences in the four-level factor session (*χ*^*2*^(5) = 29.85, *p* < 0.001), Greenhouse-Geisser corrected values are reported (*F*_(1.37, 21.96)_ = 28.21, *p* < 0.001, η_p_^2^ = 0.64). Participants gradually became faster at responding across sessions, with significant differences between session 1 (*M* = 647 ms, *SE* = 57 ms) and session 2 (*M* = 583 ms, *SE* = 48 ms; *p* < 0.001), session 2 and session 3 (*M* = 552 ms, *SE* = 48 ms; *p* = 0.006) as well as session 3 and session 4 (*M* = 533 ms, *SE* = 44 ms; *p* = 0.017). There was also a main effect of trial type (*F*_(1,16)_ = 27.46, *p* < 0.001, η_p_^2^ = 0.63). Participants responded faster in Rep trials (*M* = 560 ms, *SE* = 68 ms) as compared to Alt trials (*M* = 598 ms, *SE* = 66 ms), showing a behavioural priming effect (Olkkonen et al., 2017). For response times no other main effects or interactions were statistically significant.

The analysis of reaction times during the scanning session revealed no significant effects. Only a tendency for a faster reaction to repetition trials (*M* = 545 ms, *SE* = 24 ms) compared to alternation trials (*M* = 562 ms, *SE* = 19 ms; *F*_(1,19)_ = 3.47, *p* = 0.078, n_p_^2^ = 0.15) could be found. Descriptive data showed that response times during the scanning session (*M* = 553 ms, *SE* = 23 ms) were comparable to those from the third and fourth training sessions.

Analyses of accuracy rates during the training blocks revealed a main effect of trial type (*F*_(1,16)_ = 4.52, *p* = .049, η_p_^2^ = 0.22) with a small performance advantage for repetition (*M* = 95.9 %, *SE* = 2.3 %) than for alternation trials (*M* = 93.8%, *SE* = 2.9%). No other main effects or interactions were statistically significant. The results of the scanning session did not show any significant effects (all p’s > .5). Still, the overall performance (*M* = 95.2%, *SE* = 1.3%) showed that participants performed the task correctly.

### 3.2 Neuroimaging Results

Peak BOLD signal amplitudes by participant and condition are displayed in Figure 2A. We performed a two-by-two repeated measures ANOVA with factors context (AB, AX) and trial type (Rep, Alt) on peak BOLD signals in the right FFA (data shown in Fig 2.). There was a significant RS effect; stimuli in Rep trials evoked smaller BOLD signals (*M* = 0.65 percent signal change, *SE* = 0.08) compared to those in Alt trials (*M* = 0.71, *SE* = 0.08; main effect of trial type, *F*_(1,17)_ = 17.65, *p* < 0.001, η_p_^2^ = 0.51).

**Figure 2.**
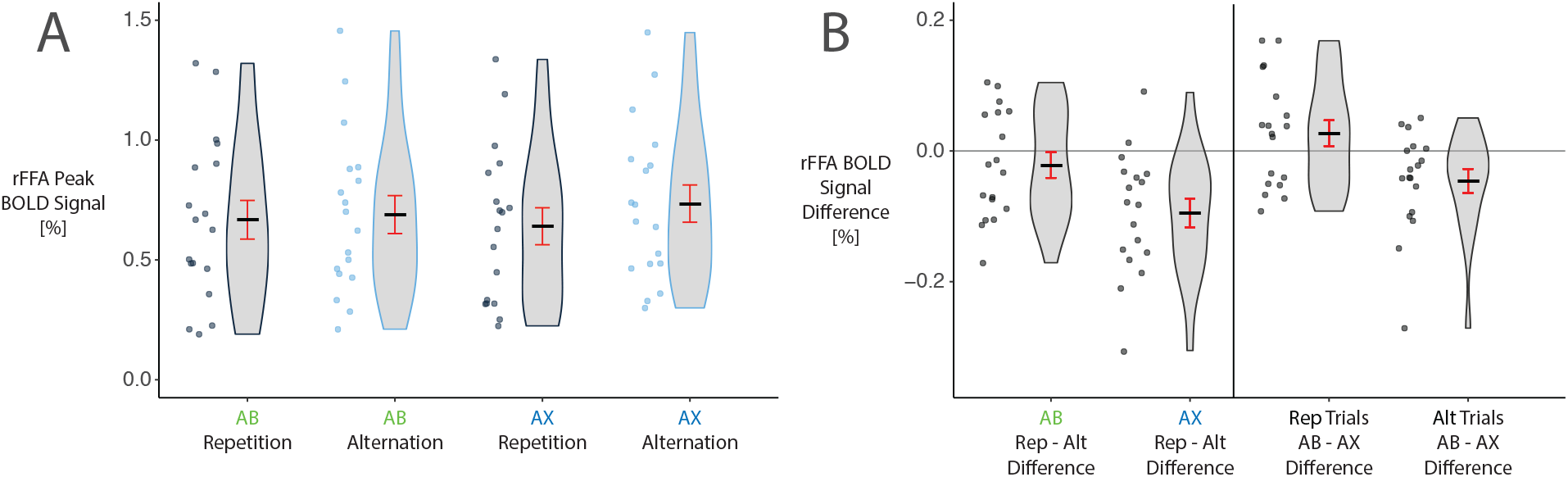
BOLD signal results for the right FFA. A) Peak BOLD signals for each Rep/Alt and AB/AX condition. Dots represent individual data points. Black lines represent group means. Error bars depict standard errors of the mean. Shaded areas depict the distributions of data for each condition. B) Repetition and context effects. Repetition effect (Rep – Alt) magnitudes for each context are shown in the left panel. Differences in BOLD signals by AB/AX context are displayed for repetition and alternation trials in the right panel.

We also found an interaction between context and trial type in the rFFA (*F*_(1,17)_ = 5.49, *p* = 0.032, η_p_^2^ = 0.24). Plotting this interaction effect revealed larger RS magnitude in the AX context (mean repetition – alternation difference = −0.095, *SE* = 0.022) as compared to the AB context (*M* = −0.022, *SE* = 0.020, shown in Fig 2B). Additionally, there appeared to be a larger magnitude effect of context on Alt trials, with AX Alt trials evoking larger BOLD signals than AB trials (mean AB – AX context effect = −0.046, *SE* = 0.018). In contrast, BOLD signals for AB and AX Rep trial differences did not differ as much (*M* = 0.027, *SE* = 0.020). Altogether, these results suggest that the extent of the observed RS largely depends on the signal magnitude of the Alt trials and this, in turn, is reduced by prior associations of S1 and S2.

As we could identify the left FFA using our localiser sequences in a subset of participants, we included this ROI in an additional analysis. We found an RS effect (main effect of trial type, *F*_(1,13)_ = 10.0, *p* 0.007, η_p_^2^ = 0.44). The interaction effect was the same pattern as found for the rFFA, but did not quite meet our statistical significance threshold (*F*_(1,13)_ = 4.36, *p* = 0.057, η_p_^2^ = 0.25).

Notably, we did not observe statistically significant RS effects in our sample in the AB context, for the right FFA (t(17) = −1.10, p = 0.287) or left FFA (t(14) = −0.23, p = 0.819, mean Rep – Alt difference = −0.004, *SE* = 0.019), whereas we did find significant RS effects in the AX context (right FFA: −4.31, p < .001; left FFA: t(14) = −3.49, p = 0.004, *M* = −0.094, *SE* = 0.027).

## 4. Discussion

To investigate the interplay between repetition and expectation effects, we presented pairs of faces which could either repeat or alternate within a trial, in two different contexts. In one context the alternating faces were chosen randomly and were therefore unpredictable, while in the other context the second face in alternating trials could be predicted after seeing the first, due to previously learned transitional rules and contingencies. In both contexts the repeated stimuli were predictable. We found repetition-related reductions of BOLD signals in the left and right FFA, consistent with a large body of work (for a review see Grill-Spector et al., 2006). More importantly, we report that responses to alternating stimuli differed markedly depending on the context; unpredictable stimulus pairs (in the AX context) evoked larger BOLD signals than those which were predictable (in the AB context). This in turn modulated the measured repetition-alternation signal differences that typically defines the measurement of RS, and even determined whether or not we found statistically-significant RS effects in our sample. Here, the point estimate of RS magnitude in the right FFA was over four times as large in the AX (*M* = 0.095) compared to AB (*M* = 0.022) contexts. Our results demonstrate that stimulus predictability effects can substantially inflate conventional measures of RS, or even mimic the effect of stimulus repetition, when predictability is not equated between repeated and alternating stimuli. While it seems unlikely that all prior reports of RS could be fully explained by effects of stimulus predictability, this effect has likely inflated repetition effect sizes in a large number of existing studies.

Our results are in line with those of Pajani et al., (2017) and (Feuerriegel et al., 2018a), who found similar effects of stimulus predictability using BOLD and ERP measures, respectively. Importantly, our design also controlled for effects of stimulus novelty across predictable and unpredictable contexts, which could have produced the patterns of effects seen in their experiments (Mur et al., 2010; Xiang and Brown, 1998). In their studies the alternating stimuli in AB-type conditions were presented many times to the participants, yet the alternating stimuli in AX-type conditions were presented much more rarely (Feuerriegel et al., 2018a) or only once in the experiment (Pajani et al., 2017). By contrast, we presented each face image in the AB and AX contexts an equal number of times, thereby replicating their findings while controlling for effects of novelty.

Notably, effects of stimulus predictability seem to be consistent across hemodynamic and electrophysiological measures, in contrast to effects of repetition probability manipulations as used in Summerfield et al. (2008) and subsequent replications. The effects of stimulus predictability seen here resemble expectations derived through statistical learning of image transition probabilities (as seen in single-cell recording measurements by Meyer & Olson, 2011), produced by the pairing of specific images, rather than more abstract expectations about whether a stimulus will repeat or not. These types of expectations appear to be qualitatively different to expectations pertaining to more abstract sequences of stimuli, and there is some evidence that these two have interacting effects on neural responses (Costa-Faidella et al., 2011; Feuerriegel et al., 2018b; Mittag et al., 2016).

Similar to the EEG study of Feuerriegel and colleagues (2018a), we observed that these context effects predominantly acted upon responses to alternating rather than repeated stimuli. This indicates that stimulus predictability selectively influenced responses to alternating stimuli, which does not modulate the underlying mechanisms of RS per se, but does influence how it is measured in commonly-used immediate repetition designs (Grill-Spector et al., 2006). This pattern of results also suggests that the violation of image-specific expectations (i.e., surprise) may underlie the observed predictability effects, and be responsible for BOLD signal increases in AX alternating trials. In our design, the likelihood of each trial type (AB-Rep, AB-Alt, AX-Rep and AX-Alt) was equated throughout the experiment. However, the relative likelihoods of the appearance of specific face images in each context were not. For example, in AB trials the S2 face could either be a repetition of S1, or a specific different face identity, with a probability ratio of 1:1. In contrast, after seeing S1 in the AX trials, an image repetition would occur 50% of the time, yet each of the 5 possible alternating face images could each appear with a probability of 10%, leading to a probability ratio of 5:1. If participants’ expectations depended on the relative appearance probabilities of specific images, then this would lead to expectations more strongly weighted toward repetitions in AX contexts, and larger surprise-related BOLD increases following AX alternating stimuli. According to this interpretation, one might also expect to see similar magnitude suppression of BOLD signals for AX repetition trials, reflecting ES, whereas we observed larger context effects for alternating trials. This may be because surprise seems to have a larger effect on neural responses than fulfilled expectations (Amado et al., 2016; Kovács and Vogels, 2014). In addition, there is evidence that effects of fulfilled expectations and surprise are diminished for repeated stimuli (reviewed in Feuerriegel et al., 2018b). So, it appears that surprise-related response enhancement in AX alternating trials may have played an important role in inflating measures of RS.

We caution that our findings should not be interpreted as that repetition effects in general are simply due to a stimulus predictability effect. Previous experiments using AB-type designs and stimulus associations have reported repetition effects (Todorovic and de Lange, 2012; Pajani et al., 2017; Feuerriegel et al., 2018a, 2019). In fact, one of the earliest mentions of RS in macaques was from the seminal study of Gross and colleagues (1979), using an AB-type design, with associated stimuli and an S1-S2 matching task.

In addition, we note that the RS effects in our study may not be strictly localized to the FFA, and may partly index inherited effects due to RS in regions early in the visual stream, such as V1, providing altered input to higher-level regions. Such ‘inherited adaptation’ effects (Kohn, 2007) have been widely documented (reviewed in Feuerriegel, 2016; Larsson, Solomon, & Kohn, 2016) and small size changes between S1 and S2 would not fully control for such effects, given the large receptive field sizes that are present in areas earlier than the FFA in the visual hierarchy. A recent optogenetic study has cast doubt on the notion that RS is locally generated in IT (Fabbrini et al., 2019), and so it remains to be seen what the magnitude of RS effects would be when controlling for both inherited adaptation and stimulus predictability. An investigation of RS in this context should aim for higher precision (i.e., more trials per participant, or a larger sample size) than in the current study and most previous studies of RS. This is because RS, which is usually a very robust effect, was not even statistically significant in the AB context in our sample, suggesting that the true magnitudes of ‘true’ RS effects may be much smaller than previously assumed.

While our findings do not provide strong evidence for or against predictive coding models that incorporate the notion of sensory precision (e.g., Auksztulewicz and Friston, 2016), it does appear that expectations can account for a proportion of repetition effects observed in many experiments. Results of recent experiments have not provided clear support for precision-based predictive coding models of RS (e.g. Amado et al., 2016; Rostalski et al., 2019; Vinken et al., 2018) and further tests of key model predictions are needed. While RS can be conceptualized as reflecting a strong prior belief towards stimuli encountered in the immediate past, it is still unclear exactly how RS fits within the broader taxonomy of expectation-related phenomena.

Our results should be interpreted with the following caveats in mind. First of all, our study used an immediate repetition design, and our results may not be generalizable to RS as measured using delayed repetition paradigms, in which a number of different intervening stimuli are presented between the first and repeated presentations of a given image (reviewed in Henson, 2016). Although predictive coding models encompass both types of repetition effects (e.g., Auksztulewicz and Friston, 2016) it is likely that these rely on different sets of neural mechanisms (Epstein et al., 2008; Weiner et al., 2010). It remains unclear whether these should be captured within a unifying framework, or if different sets of underlying mechanisms produce similar effects in each type of repetition design.

Second, we did not find differences in mean RTs and accuracy scores across AB and AX conditions, despite extensive training and exposure to the stimulus pairings. This is despite our findings of stimulus repetition effects on RTs. Because of this, it is unclear whether the predictability effects found in our neuroimaging results were actually used for decision making during the task. Although validly-cued expectancies for certain stimuli have led to faster responses in previous studies (Hall et al., 2018; Mulder et al., 2012), these designs have typically conflated expectations to see a certain stimulus with preparation of motor actions corresponding to that stimulus (Gold and Stocker, 2017). In addition, recent findings have cast doubt on the idea that contextual expectations affect those sensory representations that are used for perceptual decision making, at least in the same trials whereby those expectations are fulfilled or violated, and when controlling for feature-selective attention (Bang and Rahnev, 2017; Rungratsameetaweemana et al., 2018). In our task there were no cued biases toward a particular button response, and participants’ expectations for how to respond were balanced across AB and AX contexts. This may be why we did not observe predictability effects on behavior.

## 5. Conclusion

We have shown that, in immediate repetition designs, an observer’s capacity to predict the image of repeated compared to unrepeated stimuli has a substantial effect on the observed magnitude of RS. While this does not necessarily mean that RS is best accounted for by predictive coding models, it does indicate that measures of repetition effects have likely been inflated due to this confound in a very large number of previous studies, including those run within our own labs. We also highlight stimulus predictability as an important, yet commonly overlooked, factor to consider when investigating the hierarchy of expectation effects implemented within the visual system.

## Acknowledgements

The authors would like to thank Carolin Altmann for providing the stimulus set. This work was supported by a grant from the Deutsche Forschungsgemeinschaft (KO3918/5-1).

